# APEX2 proximity proteomics resolves flagellum subdomains and identifies flagellum tip-specific proteins in *Trypanosoma bruce*i

**DOI:** 10.1101/2020.03.09.984815

**Authors:** Daniel E. Vélez-Ramírez, Michelle M. Shimogawa, Sunayan Ray, Andrew Lopez, Shima Rayatpisheh, Gerasimos Langousis, Marcus Gallagher-Jones, Samuel Dean, James A. Wohlschlegel, Kent L. Hill

## Abstract

*Trypanosoma brucei* is the protozoan parasite responsible for sleeping sickness, a lethal vector-borne disease. *T. brucei* has a single flagellum that plays critical roles in parasite biology, transmission and pathogenesis. An emerging concept in flagellum biology is that the organelle is organized into subdomains, each having specialized composition and function. Overall flagellum proteome has been well-studied, but a critical gap in knowledge is the protein composition of individual flagellum subdomains. We have therefore used APEX-based proximity proteomics to examine protein composition of *T. brucei* flagellum subdomains. To assess effectiveness of APEX-based proximity labeling, we fused APEX2 to the DRC1 subunit of the nexin-dynein regulatory complex, an axonemal complex distributed along the flagellum. We found that DRC1-APEX2 directs flagellum-specific biotinylation and purification of biotinylated proteins yields a DRC1 “proximity proteome” showing good overlap with proteomes obtained from purified axonemes. We next employed APEX2 fused to a flagellar membrane protein that is restricted to the flagellum tip, adenylate cyclase 1 (AC1), or a flagellar membrane protein that is excluded from the flagellum tip, FS179. Principal component analysis demonstrated the pools of biotinylated proteins in AC1-APEX2 and FS179-APEX2 samples are distinguished from each other. Comparing proteins in these two pools allowed us to identify an AC1 proximity proteome that is enriched for flagellum tip proteins and includes several proteins involved in signal transduction. Our combined results demonstrate that APEX2-based proximity proteomics is effective in *T. brucei* and can be used to resolve proteome composition of flagellum subdomains that cannot themselves be readily purified.

**IMPORTANCE:** Sleeping sickness is a neglected tropical disease, caused by the protozoan parasite *Trypanosoma brucei*. The disease disrupts the sleep-wake cycle, leading to coma and death if left untreated. *T. brucei* motility, transmission, and virulence depend on its flagellum (aka cilium), which consists of several different specialized subdomains. Given the essential and multifunctional role of the *T. brucei* flagellum, there is need of approaches that enable proteomic analysis of individual subdomains. Our work establishes that APEX2 proximity labeling can, indeed, be implemented in the biochemical environment of *T. brucei*, and has allowed identification of proximity proteomes for different subdomains. This capacity opens the possibility to study the composition and function of other compartments. We further expect that this approach may be extended to other eukaryotic pathogens, and will enhance the utility of *T. brucei* as a model organism to study ciliopathies, heritable human diseases in which cilia function is impaired.

## INTRODUCTION

*Trypanosoma brucei* is a flagellated parasite that is transmitted between mammalian hosts by a hematophagous vector, the tsetse fly, and is of medical relevance as the causative agent of sleeping sickness in humans [1]. *T. brucei* also presents a substantial economic burden in endemic regions due to infection of livestock, causing an estimated loss of over 1 billion USD/year [2]. As such, *T. brucei* is considered both cause and consequence of poverty in some of the poorest regions in the world. *T. brucei* also provides an excellent model system for understanding the cell and molecular biology of related flagellates *T. cruzi* and *Leishmania spp*., which together present a tremendous health burden across the globe.

*T. brucei* has a single flagellum that is essential for parasite viability, infection and transmission [3–5]. The flagellum drives parasite motility, which is necessary for infection of the mammalian host [4], and for transmission by the tsetse fly [5]. In addition to its canonical function in motility, the flagellum plays important roles in cell division and morphogenesis [6–8] and mediates direct interaction with host tissues [9]. Moreover, recent work has demonstrated that the trypanosome flagellum is the site of signaling pathways that control the parasite’s response to external signals and are required for transmission and virulence [4, 10–15].

The trypanosome flagellum is built on a canonical “9 + 2” axoneme that originates at the basal body in the cytoplasm near the posterior end of the cell [3]. Triplet microtubules of the basal body extend to become doublets in the transition zone, which marks the boundary between the basal body and 9 + 2 axoneme [16]. The axoneme exits the cytoplasm through a specialized invagination of the plasma membrane, termed the “flagellar pocket” (FP) [17]. As it emerges from the flagellar pocket, the axoneme is attached to an additional filament, termed the “paraflagellar rod” (PFR) [18], that extends alongside the axoneme to the anterior end of the cell. The axoneme and PFR remain surrounded by cell membrane and this entire structure is laterally attached to the cell body along its length, except for a small region at the distal tip that extends beyond the cell’s anterior end [19]. Lateral flagellum attachment is mediated by proteins in the flagellum and cell body that hold the flagellum and plasma membranes in tight apposition, constituting a specialized “flagellar attachment zone” (FAZ) that extends from the flagellar pocket to the anterior end of the cell [19, 20].

As in other flagellated eukaryotes, the trypanosome flagellar apparatus (Fig. 1) can be subdivided into multiple subdomains, each having specialized function and protein composition. The FP, for example, demarcates the boundary between the flagellar membrane and cell membrane. In *T. brucei*, the FP is the sole site for endocytosis and secretion, thus presenting a critical portal for host-parasite interaction [17]. The basal body functions in flagellum duplication, segregation and axoneme assembly [21]. The region encompassing the transition zone lies between the cytoplasm and flagellar compartment and includes proteins that control access into and out of the flagellum [13, 16]. The 9 + 2 axoneme is the engine of motility [7], while the PFR in trypanosomes is considered to physically influence flagellum beating and to serve as a scaffold for assembly of signaling and regulatory proteins [22–25]. The FAZ is a trypanosome-specific structure and is critical for parasite motility and cell morphogenesis [26]. The flagellum tip marks the site of cleavage furrow initiation during cytokinesis [27] and mediates attachment to the tsetse fly salivary gland epithelium, which is crucial for development into human infectious parasites [28]. The flagellum tip is also the site of signaling proteins that function in cell-cell communication and signaling [11, 29].

**Figure 1.**
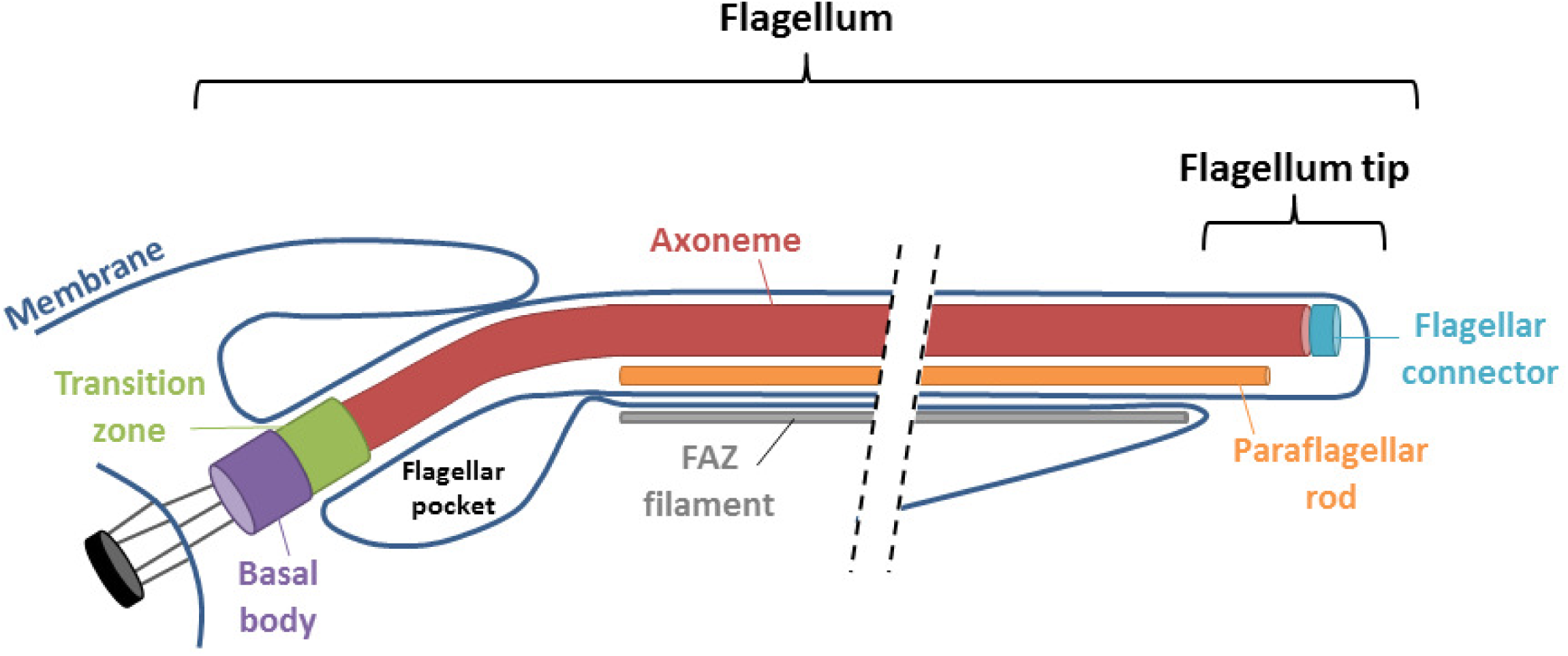
Schematic diagram of major flagellum substructures. Schematic depicts emergence of the flagellum from the basal body near the cell’s posterior end (left), extending to the cell’s anterior end (right). Major substructures are labeled.

Given the essential and multifunctional roles of the trypanosome flagellum, much effort has been made to define the protein composition of the organelle. However, although we know a lot about protein composition of the flagellum as a whole, a critical knowledge gap is the protein composition of individual flagellum subdomains. Classical proteomics approaches typically require purification of the flagellum, or subfractions of the flagellum [16, 30–34]. This approach is very useful and has been used to determine proteomes of the transition zone [16] and axoneme tip [31], but it can be cumbersome, is dependent on quality of the purified fraction and cannot resolve subdomains that are not part of a specific structure that can be purified.

To overcome limitations of conventional flagellum proteomics approaches, we have adapted APEX2 proximity labeling [35] for use in *T. brucei*. APEX2 is an engineered monomeric ascorbate peroxidase that converts biotin-phenol into a short-lived biotin radical that is highly reactive. The biotin-phenol radical interacts with nearby proteins, resulting in covalent attachment of a biotin tag. Biotinylated proteins can be affinity-purified with streptavidin and identified by shotgun proteomics, allowing for facile identification of proteins within a specific subcellular location from a complex and largely unfractionated sample [35, 36]. Here, we report successful implementation of APEX2 proximity labeling in *T. brucei*, to define the proximity proteome of flagellar proteins that are either distributed along the axoneme or restricted to the tip of the flagellum membrane. Our results establish APEX-based proximity proteomics as a powerful tool for *T. brucei*, demonstrate the approach can resolve flagellum subdomains that are not separated by a physical boundary and support the idea that the flagellum tip subdomain is specialized for cell signaling.

## RESULTS

To evaluate APEX2 labeling in *T. brucei*, we selected an axonemal protein as bait, because the axoneme is a well-defined cellular component whose protein composition in *T. brucei* has been examined in prior studies [16, 30, 31, 34]. We selected the DRC1 subunit of the nexin-dynein regulatory complex (NDRC) [37], because this protein has a defined localization along the axoneme and its position relative to major axonemal substructures, e.g. microtubule doublets, radial spokes and dynein arms, is known [38]. Having selected DRC1 as bait, we used *in situ* gene tagging [39] to generate cell lines expressing DRC1 fused to a C-terminal APEX2 tag that includes an HA-tag following the APEX2 tag, referred to as “DRC1-APEX2”. Expression of DRC1-APEX2 was demonstrated in Western blots of whole cell lysates (Fig. 2A). Extraction with non-ionic detergent leaves the axoneme intact in a detergent-insoluble cytoskeleton fraction that can be isolated from detergent-soluble proteins by centrifugation [40]. We found that DRC1-APEX2 fractionates almost completely with the detergent-insoluble cytoskeleton as expected for an NDRC protein (Fig. 2A). Growth curves demonstrated that expression of DRC1-APEX2 does not affect growth of *T. brucei in vitro* (Supplemental Fig. S1).

**Figure 2.**
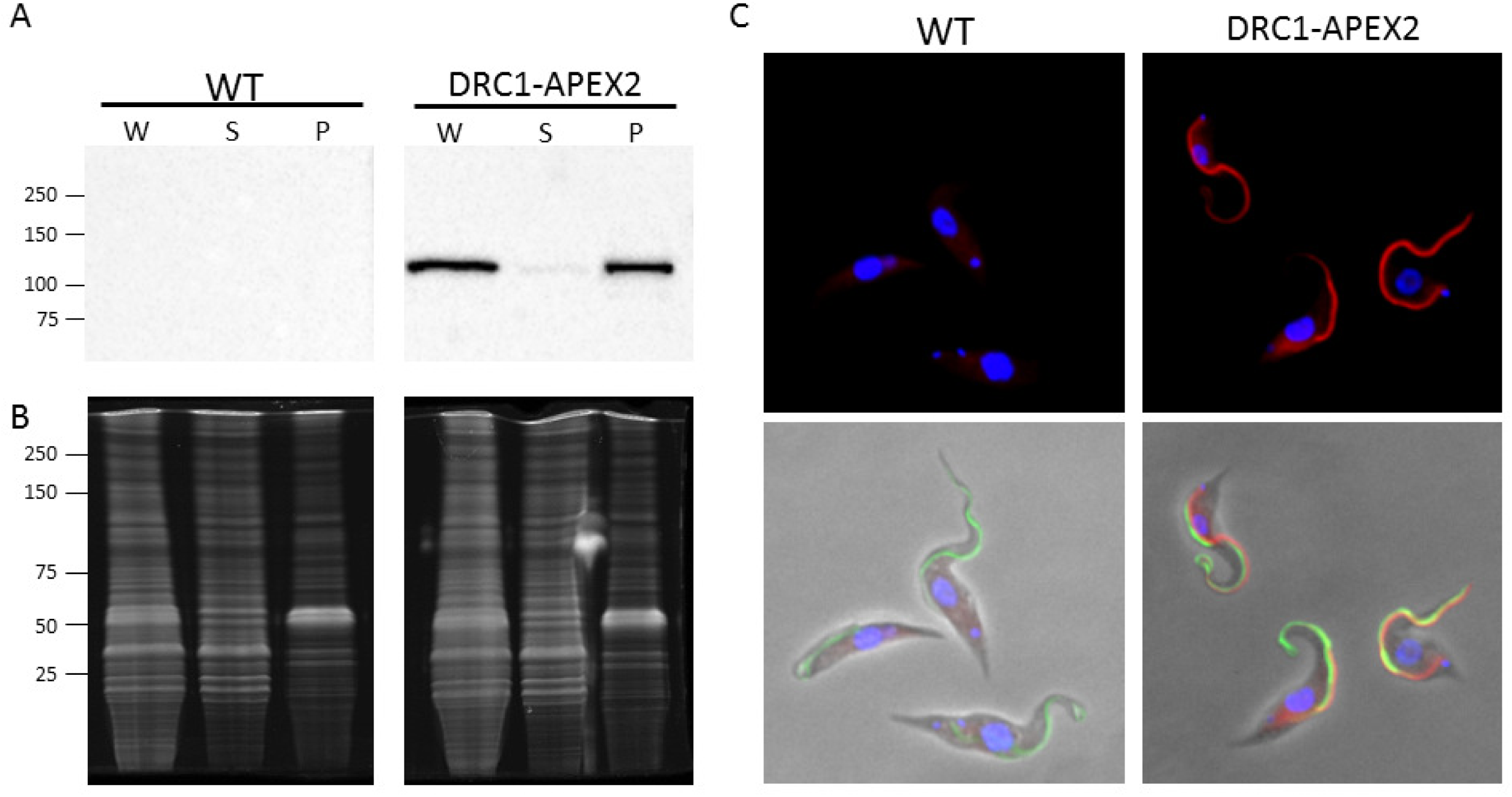
APEX2 directs organelle-specific biotinylation in *T. brucei*. Panel A. Western blot of whole cell lysate (W), NP40-extracted supernatant (S) and pellet (P) samples from WT and DRC1-APEX2-expressing cells. Samples were probed with anti-HA antibody. Panel B. Samples as in A were stained with SYPRO Ruby to assess loading. Panel C. WT and DRC1-APEX2 cells were examined by immunofluorescence with anti-PFR antibody (Alexa 488, green), Streptavidin (Alexa 594, red) and DAPI.

We next asked if APEX2 is functional within the biochemical environment of the *T. brucei* cell. Cells expressing DRC1-APEX2 were incubated with biotin-phenol, which was then activated with brief H_2_O_2_ treatment followed by quenching with trolox and L-ascorbate. Cells were then probed with streptavidin-Alexa 594 and subjected to fluorescence microscopy to assess biotinylation. As shown in Figure 2C, we observed APEX2-dependent biotinylation and this was highly enriched in the flagellum. There was some background staining in the cytoplasm, as revealed by parallel analysis of parental cells lacking the APEX2-tagged protein (Fig. 2C), but flagellum staining was only observed in cells expressing DRC1-APEX2. In the proximal region of the flagellum, the streptavidin signal extended further than the PFR (Fig. 2C), indicating streptavidin labeling is on the axoneme. Therefore, DRC1-APEX2 directs specific biotinylation in the flagellum.

Having established that DRC1-APEX2 directs flagellum-specific biotinylation, we used shotgun proteomics to identify biotinylated proteins. Samples were extracted with non-ionic detergent and separated into detergent-soluble supernatant and detergent-insoluble pellet fractions. Biotinylated proteins in each fraction were then isolated using streptavidin purification and subjected to shotgun proteomics for protein identification. WT and DRC1-APEX2 cells were processed in parallel (Fig. 3A). Our focus was the detergent-insoluble pellet because this fraction includes the axoneme and PFR, and the protein composition of these structures has been characterized [19, 25, 30, 34, 41]. The pellet fraction also includes non-axonemal structures such as the basal body, tripartite attachment complex, cytoplasmic FAZ filament and subpellicular cytoskeleton, thus enabling us to test for enrichment of axonemal proteins. The analysis was done using three independent biological replicates. In one case, the sample was split into two aliquots and shotgun proteomics was done on both in parallel, giving a total of four replicates each for DRC1-APEX2 and WT pellet samples.

**Figure 3.**
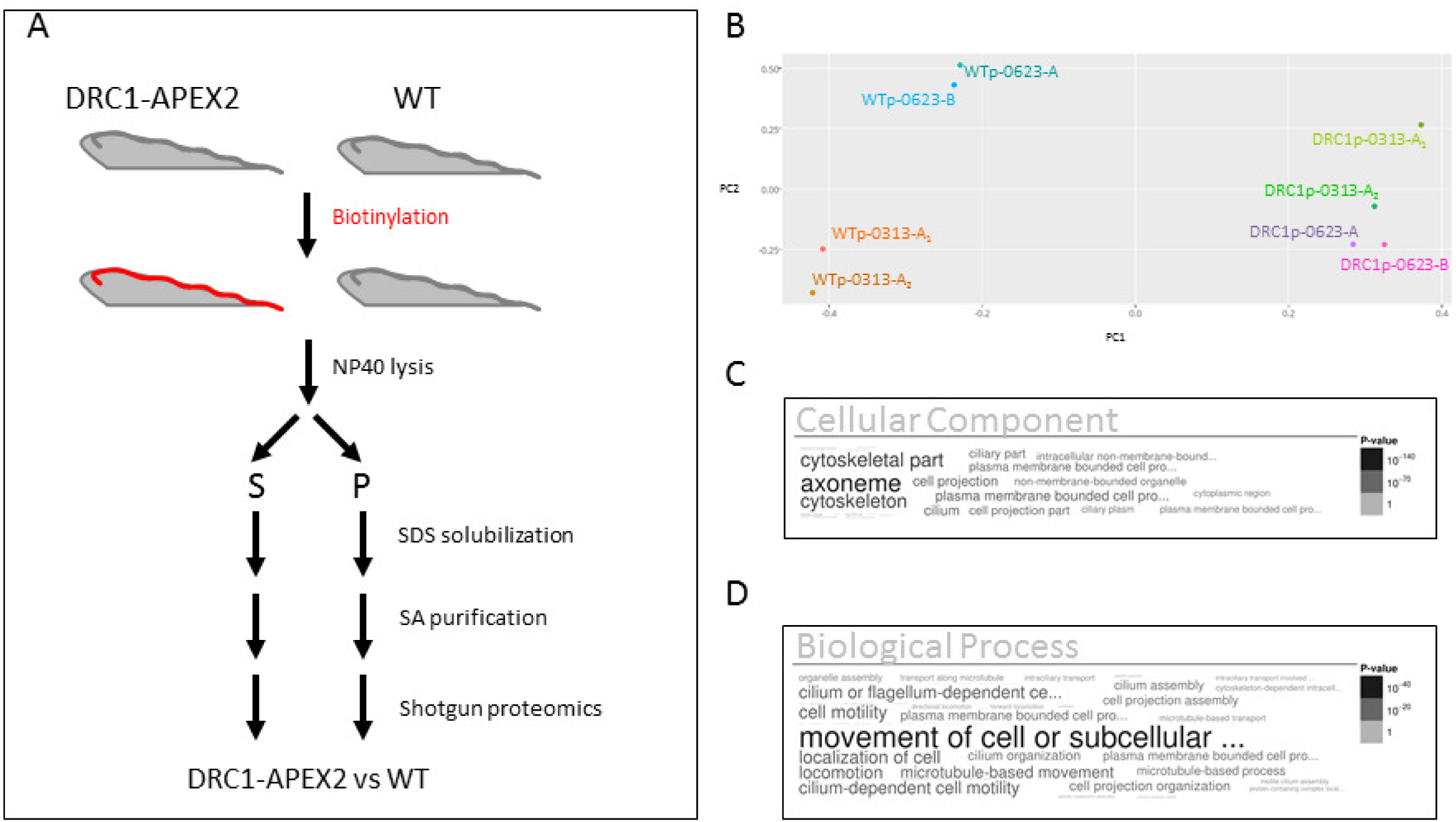
DRC1-APEX2 proximity proteome is enrichened for flagellar proteins. Panel A. Scheme used to identify biotinylated proteins from WT and DRC1-APEX2 expressing *T. brucei* cells. Panel B. Principal Component Analysis of proteins identified in pellet fractions from WT cells (WTp) and DRC1-APEX2 cells (DRC1p). The experiment was done using three independent biological replicates (0313, 0623-A and 0623-B), and the 0313 protein sample was split into two aliquots and shotgun proteomics done on each in parallel (0313-A1 and 0313-A2). Panels C and D. Word clouds showing GO analysis of the DRC1p proximity proteome. Size and shading of text reflects the p-value according to the scale shown.

Principal component analysis demonstrated that the biotinylated protein profile of the DRC1-APEX2 pellet samples (DRC1p) was distinct from that of WT samples (WTp) processed in parallel (Fig. 3B). We detected bona fide axonemal proteins, including DRC1 and axonemal dynein subunits in some WT replicates, but these were enriched in DRC1-APEX2 samples relative to WT. We therefore assembled a “DRC1p proximity proteome” that included only proteins meeting the following three criteria: i) detected in all four replicates of the DRC1p sample, ii) had a normalized spectrum count of two or more, and iii) were enriched in the DRC1p versus WTp sample. This yielded a DRC1p proximity proteome of 697 proteins (Supplemental Table S1). Human homologues were identified for 372 proteins in the DRC1p proximity proteome and among these, 38 are linked to human diseases that have been connected to cilium defects (Table 1).

**Table 1.**
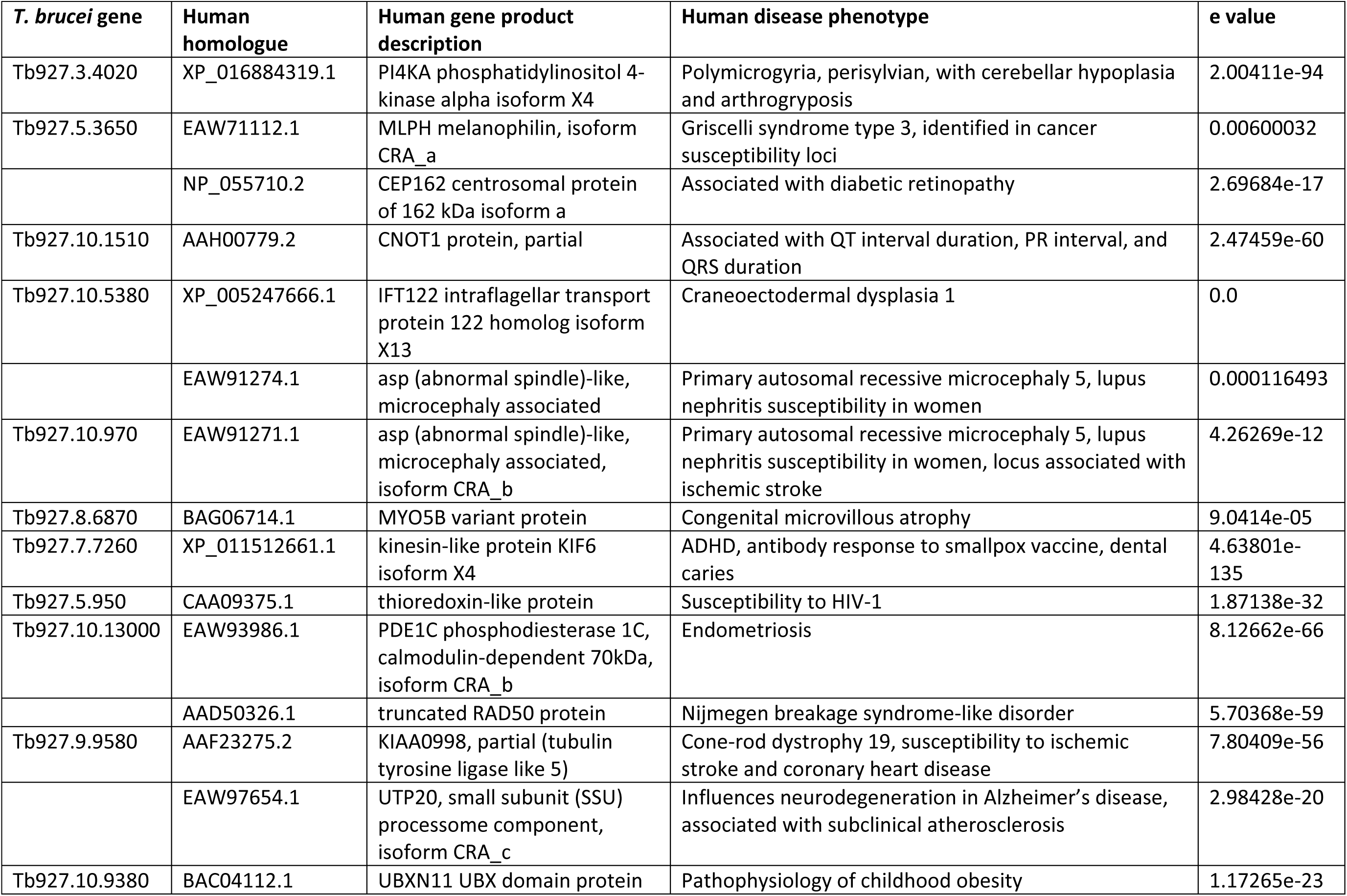
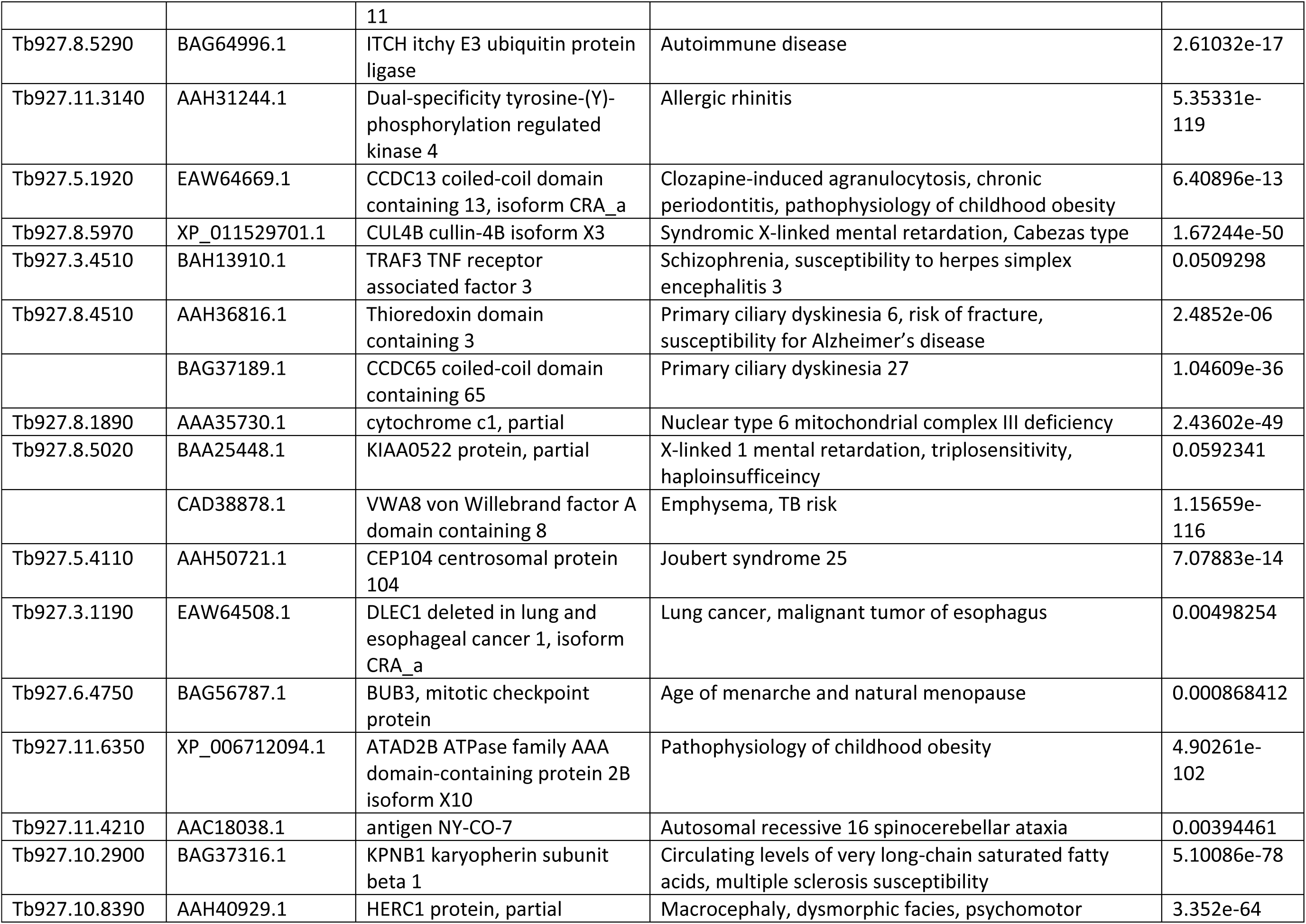
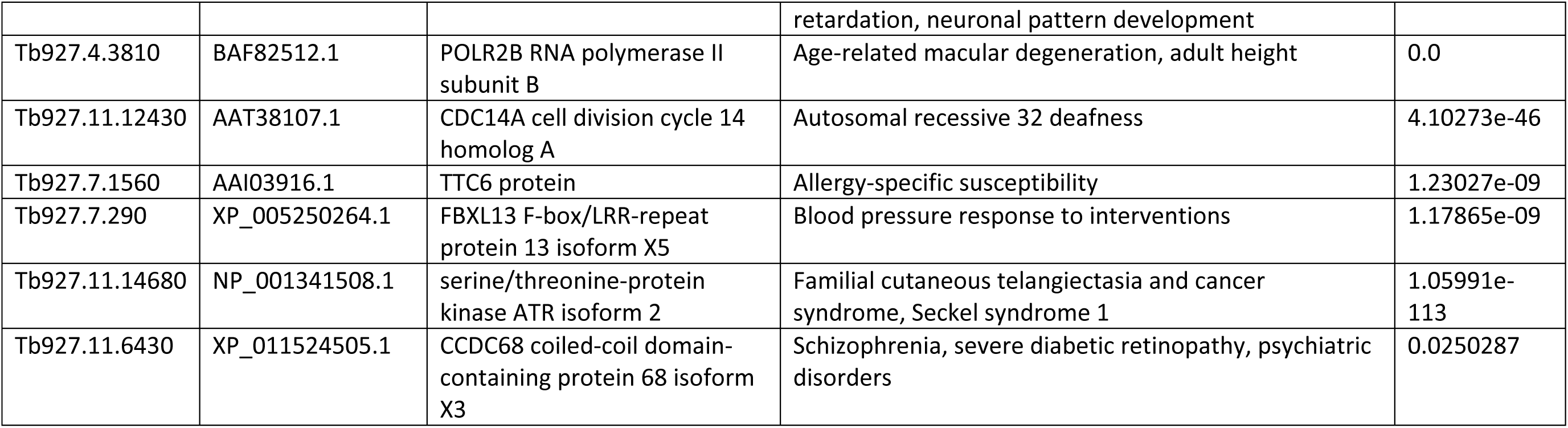
Human homologues with disease association

To evaluate whether APEX proximity labeling was effective in identifying flagellar proteins, we used Gene Ontology (GO) analysis [42], comparison to prior *T. brucei* flagellar proteomes [30, 32, 34] and independent tests of localization [43]. GO analysis demonstrated significant enrichment of flagellar proteins in the DRC1p proximity proteome compared to the genome as a whole (Fig. 3C and D). As discussed above, our efforts were focused on the pellet fraction. We did however, complete GO analysis on the detergent-soluble “DRC1s proximity proteome”, which also showed significant enrichment of flagellar proteins, as well as signaling proteins (Supplemental Fig. S2).

When compared with prior proteomic analyses of *T. brucei* flagella, the DRC1p proximity proteome encompassed a larger fraction (45%) of the flagellum skeleton proteome [30] than of the intact flagellum proteome (36%) [34], perhaps due to the fact that the latter includes detergent-soluble proteins, which are not expected in the DRC1p proximity proteome. As anticipated, minimal overlap was observed with the flagellum surface plus matrix proteome [32], which includes only detergent-soluble proteins.

TrypTag localization data [43] were available for 677 proteins in the DRC1p proximity proteome and 509 of these (75%) are annotated as having TrypTag localization that includes one or more flagellum structures. The DRC1p proximity proteome includes 350 proteins that were not identified in prior proteomic analyses of the *T. brucei* flagellum or axoneme fragments [16, 25, 30–32, 34] (Supplemental Table S1A). TrypTag localization data were available for 337 of these and 241 (71%) are annotated as having a TrypTag localization to one or more flagellum structures. In some cases, localization was specific to flagellum structures, while in others the protein showed multiple locations. This finding supports the idea that many of these 350 proteins are bona fide flagellar proteins despite going undetected in earlier flagellum proteome studies. The combined results demonstrate that APEX2 proximity labeling is functional in *T. brucei* and enables identification of flagellar proteins without the need to purify the flagellum. The data also indicate protein composition of the *T. brucei* flagellum is more complex than indicated by earlier studies alone.

APEX labeling readily distinguishes proteins in close proximity but separated by a membrane [36] and this is evidenced in our data when considering protein components of the FAZ [19]. Proteins on the flagellar side of the FAZ are substantially enriched in the DRC1-APEX2 sample, whereas proteins on the cell body side of the FAZ are not (Supplemental Fig. S3). Furthermore, the short half-life of the biotin-phenol radical [35] means that APEX labeling can resolve proteins separated by distance even in the absence of a membrane boundary. As discussed above and shown previously [44], APEX resolves flagellar versus cytoplasmic proteins despite these two compartments being contiguous. Within the DRC1p proximity proteome, we noted that proteins distributed similarly to DRC1 along the axoneme were well-represented, while proteins restricted to the distal or proximal end of the flagellum were less represented (Fig. 4 and Supplemental Table S2). While total abundance may contribute to this result, it nonetheless suggested that beyond flagellum versus cytoplasm, APEX labeling might also be able to resolve proteins from different subdomains within the flagellum. We were particularly interested in the flagellum tip because of its importance in trypanosomes and other organisms for signal transduction [10, 11, 45, 46], flagellum length regulation [47–51] and interaction with host tissues [9].

**Figure 4.**
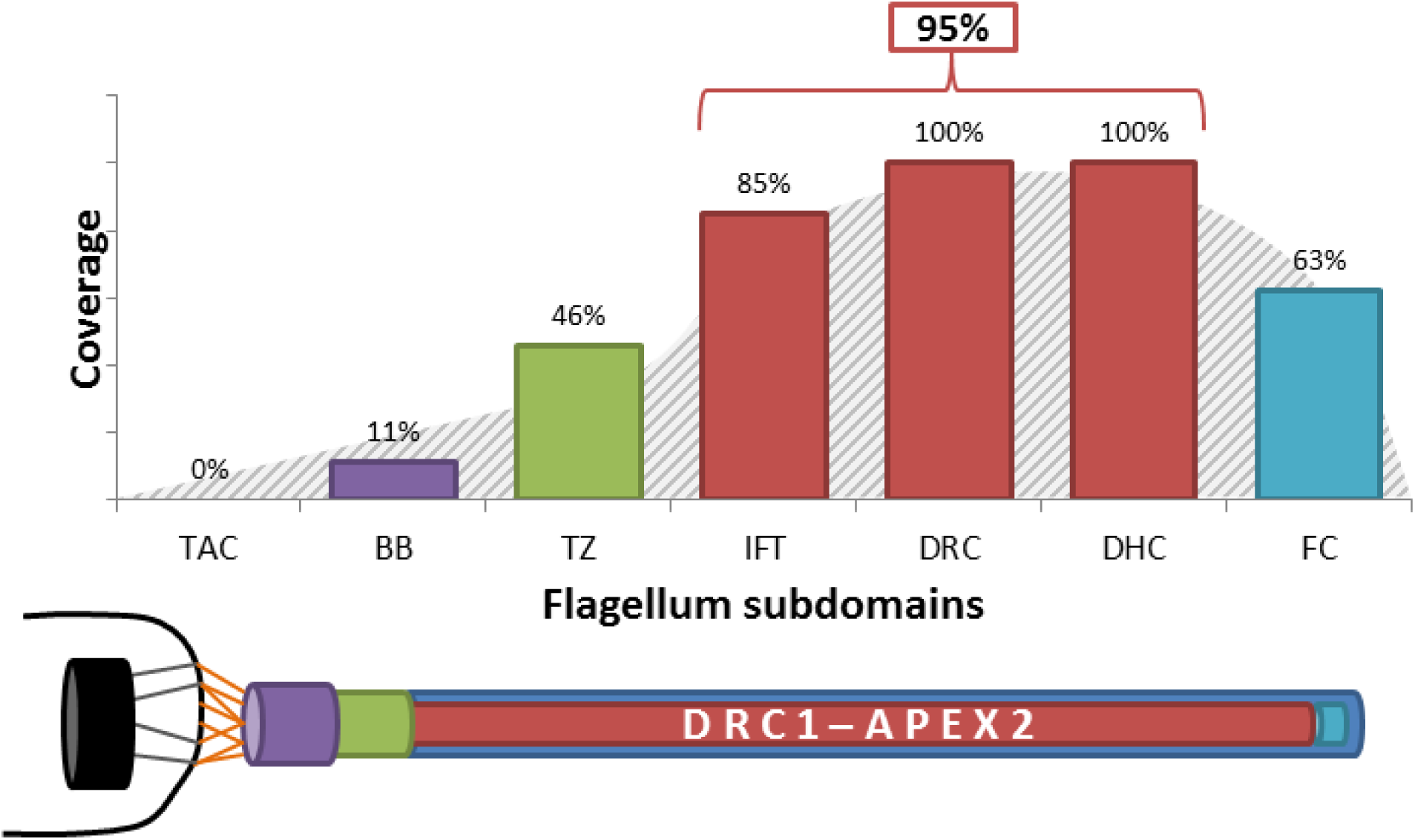
Spatial distribution of proteins identified in the DRC1p proximity proteome. Bars in the histogram indicate the percentage of proteins from each of the indicated complexes that were identified in the DRC1p proximity proteome. Schematic below the histogram illustrates the relative distribution of the complexes indicated in the histogram, with the mitochondrion and kinetoplast indicated in black at left.

We generated APEX2-tagged versions of flagellar proteins that are tip-specific, AC1 [29], or tip-excluded, FS179 [32]. Expression of either AC1-APEX2 or FS179-APEX2 did not affect *T. brucei* doubling time (Supplemental Fig. S1) and both tagged proteins fractionated in the detergent-soluble fraction as expected. To assess biotinylation, AC1-APEX2 and FS179-APEX2-expressing cells were incubated with biotin-phenol, activated with H_2_O_2_, then quenched, probed with streptavidin-Alexa 594 and examined by fluorescence microscopy (Fig. 5A-D). The signal in AC1-APEX2 expressors was enriched at the flagellum tip, while the signal in FS179-APEX2 expressors was distributed along the flagellum but lacking or diminished at the flagellum tip (Fig. 5A-D). Samples were solubilized with detergent and centrifuged to remove insoluble material. Biotinylated proteins were isolated from the soluble fraction by streptavidin affinity purification and then identified by shotgun proteomics. As negative controls, samples from WT cells without an APEX2 tag and cells expressing AC1Δ45-APEX2, an AC1 truncation that lacks the C-terminal 45aa and is localized to the cytoplasm instead of the flagellum [29], were processed in parallel (Fig. 5E).

**Figure 5.**
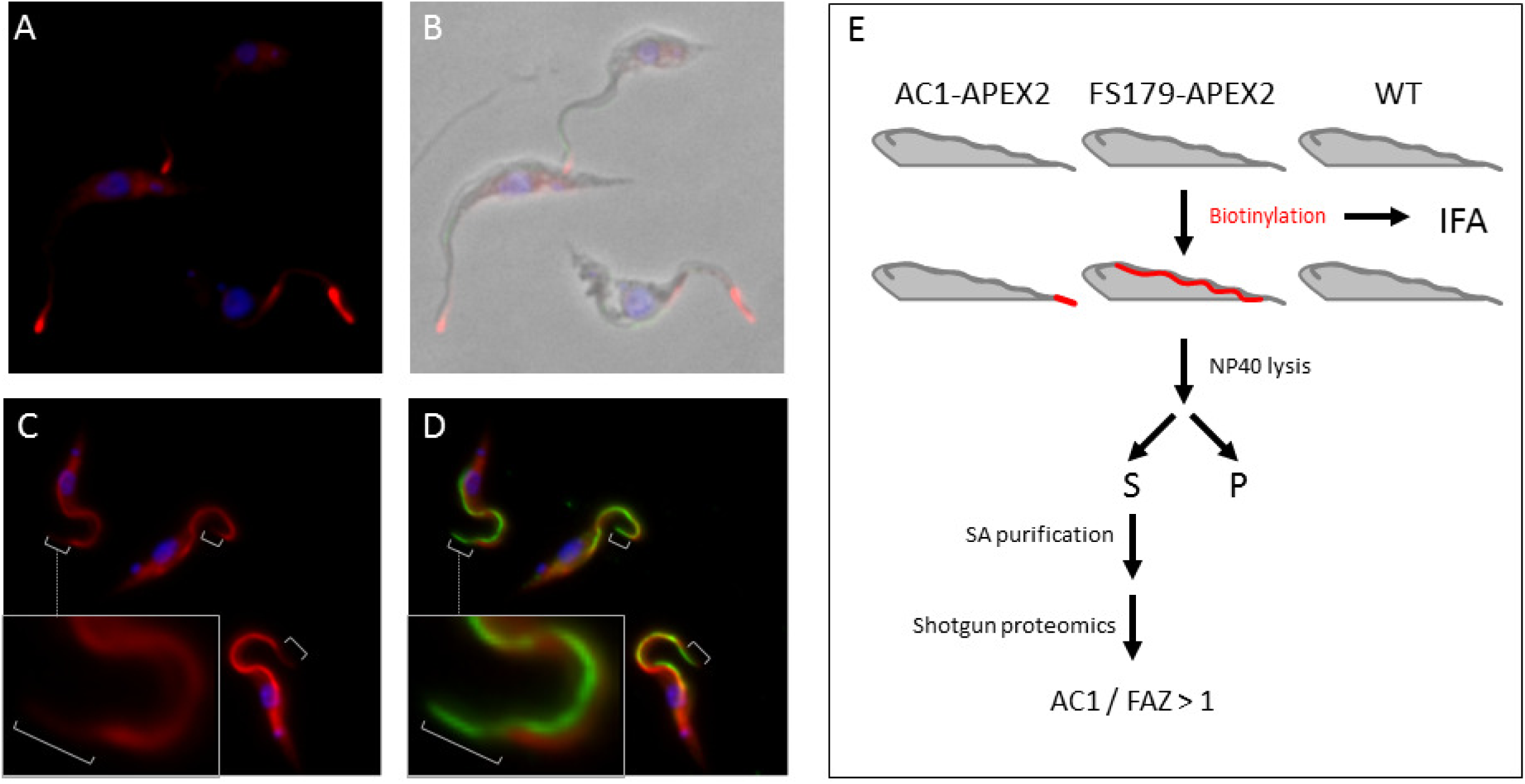
APEX2-labeling resolves flagellum sub-domains. Panels A, B. AC1-APEX2 cells were fixed and examined by fluorescence microscopy after staining with Streptavidin (red) and DAPI (blue). Panels C, D. FS179-APEX2 cells were examined by immunofluorescence with anti-PFR antibody (Alexa 488, green), Streptavidin (Alexa 594, red) and DAPI (blue). White brackets indicate the distal region of the flagellum that is not labelled by streptavidin. Box shows zoomed in version of the cell in the middle of the field. Panel E. Scheme used to identify biotinylated proteins from the indicated cell lines (WT, AC-APEX2 and FS179-APEX2).

Principal component analysis demonstrated that the biotinylated protein profile of AC1-APEX2 detergent-soluble (AC1s) samples was readily distinguished from that of FS179-APEX2 (FS179s) and WT (2913s) controls (Fig. 6A). Therefore, APEX2 proximity proteomics was able to distinguish protein composition of the tip versus FAZ subdomains within the flagellum, even though they have no physical barrier between them.

**Figure 6.**
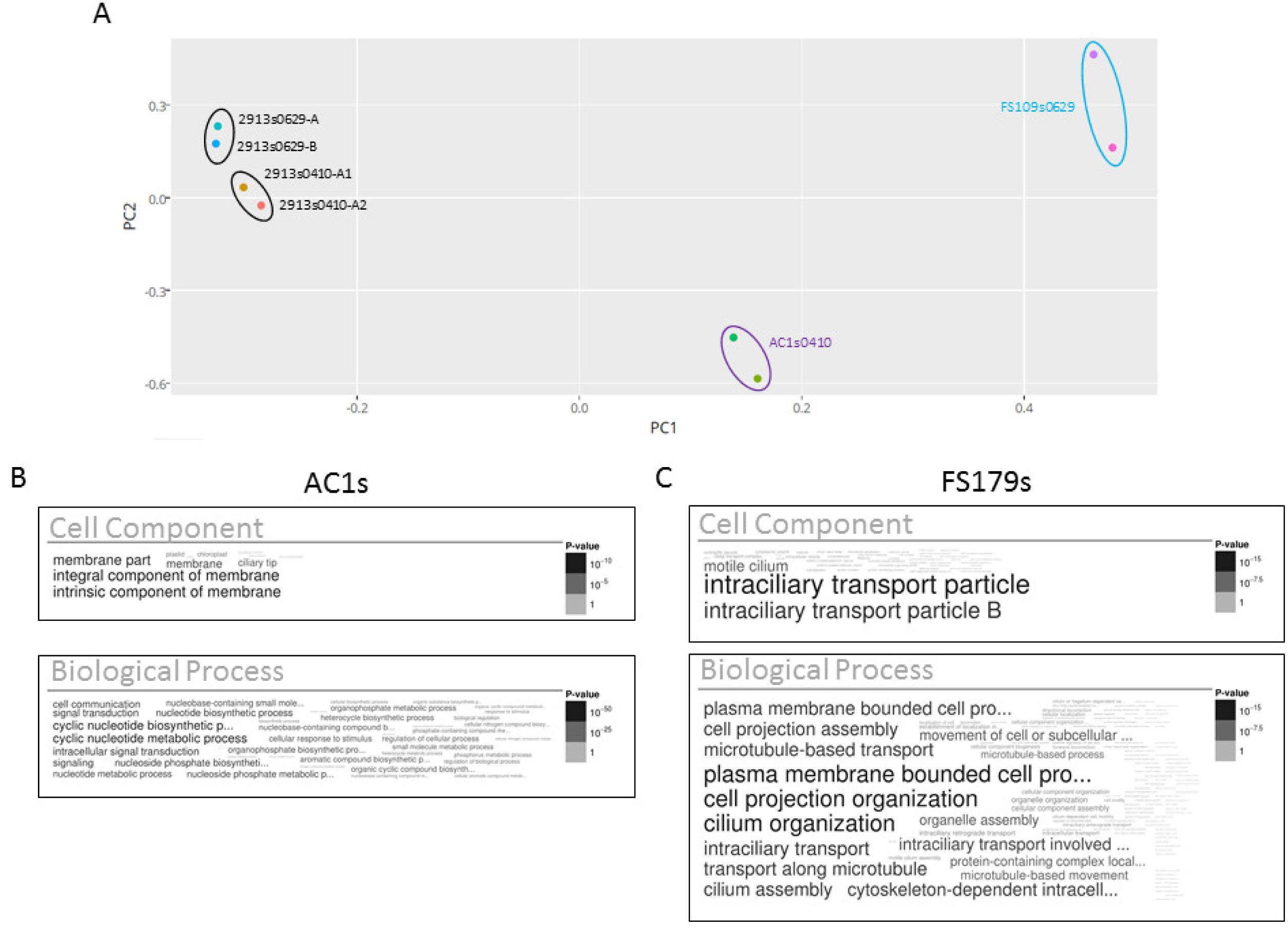
APEX2 proximity proteomics differentiates protein composition of distinct flagellum subdomains. Panel A. Principal Component Analysis of proteins identified in supernatant fractions from WT (2913s), AC1-APEX2 (AC1s) and FS179-APEX2 (FS179s) cells. For 2913 controls, three independent experiments are shown (0410, 0629-A and 0629-B) and for one experiment the sample was split into two aliquots (0410-A1 and 0410-A2) that were subjected to shotgun proteomics in parallel. For FS179s, two independent experiments are shown (0629-A, 0629-B). For AC1s, one sample was split into two aliquots that were subjected to shotgun proteomics in parallel (0410-A1, 0410-A2). Panel B. Word cloud representing GO analysis for Cellular Component and Biological Process of the proteins identified in the AC1 proximity proteome. Panel C. Word cloud representing GO analysis for Cellular Component and Biological Process of the proteins identified in the FS179 proximity proteome.

To define an “AC1s proximity proteome”, we compared the biotinylated protein profile of the AC1s sample to that of 2913s and AC1Δ45s samples processed in parallel. We used known flagellum tip proteins (Supplemental Table S3) to set the enrichment threshold for inclusion in the AC1s proximity proteome. Finally, we included only those proteins that were enriched in AC1s versus FS179s and DRC1s. This yielded a final AC1s proximity proteome of 48 proteins, including 27 adenylate cyclases and 21 additional proteins (Table 2). Extensive sequence identity among adenylate cyclases poses challenges for distinguishing between some isoforms, so the number of 27 might be an overestimate. Nonetheless, adenylate cyclases represent the majority of proteins identified. We performed a parallel analysis to define an “FS179s proximity proteome”, in this case comparing to 2913s and AC1Δ45s samples as negative controls and using intraflagellar transport (IFT) proteins [52] to set the enrichment threshold for inclusion. Gene ontology analysis showed enrichment of flagellar proteins and signaling proteins in the AC1s proximity proteome, while the FS179s proximity proteome is enriched for flagellar proteins, but not signaling proteins (Fig. 6B and C).

**Table 2.** AC1s proximity proteome.

| TriTryp GeneID | TriTryp Annotation | Localization* | Source for location data | AC1s/FS179s avg ratio<sup>&</sup> |
| --- | --- | --- | --- | --- |
| Tb927.2.5860 | Hypothetical | **Tip** | TrypTag<sup>1</sup> | 15.88 |
| Tb927.2.5870 | Hypothetical | **Tip** | TrypTag<sup>1</sup> + this work | 10.71 |
| Tb927.4.1500 | RNA editing associated helicase 2 | **Not Flagellar** | TrypTag<sup>1</sup> | 7.45 |
| Tb927.4.4400 | Hypothetical | **Flagellum, distal points + Endocytic** | TrypTag<sup>1</sup> | 5.18 |
| Tb927.2.5760 | Flagellar Member 8 | **Tip** | 2 | 5.03 |
| Tb927.11.14410 | Ankyrin repeats | **Not Flagellar** | TrypTag<sup>1</sup> | 3.89 |
| Tb927.1.2120 | Calpain-like protein CALP1.3 | **Tip** | 3 | 3.60 |
| Tb927.11.17040 | AC1<sup>#</sup> | **Tip** | 4 | 3.37 |
| Tb927.7.4060 | cysteine peptidase, Clan CA, family C2 | **Flagellum, Tip-enriched + Cytoplasm** | TrypTag<sup>1+3</sup> | 3.02 |
| Tb927.9.7540 | cysteine peptidase, Clan CA, family C2, putative | **Flagellum, Tip-enriched** | TrypTag<sup>1</sup> + this work | 2.98 |
| Tb927.7.4070 | cysteine peptidase, Clan CA, family C2 | **Tip + Cytoplasm** | 3 | 2.86 |
| Tb927.7.5340 | cAMP response protein 3 | **Flagellum, Tip-enriched + Cytoplasm** | TrypTag + this Work | 2.41 |
| Tb927.10.15700 | Hypothetical | **NA** |  | 2.11 |
| Tb927.4.4220 | small GTP-binding rab protein | **NA** |  | 2.11 |
| Tb927.10.9380 | SEP domain/UBX domain containing protein | **New Flagellum-distal points** | TrypTag<sup>1</sup> | 1.86 |
| Tb927.5.2090 | kinesin | **Flagellum + Cytoplasm** | TrypTag<sup>1</sup> | 1.74 |
| Tb927.6.4710 | calmodulin | **Flagellum** | TrypTag<sup>1</sup> | 1.68 |
| Tb927.3.4640 | VIT family | **Not Flagellar** | TrypTag<sup>1</sup> | 1.50 |
| Tb927.7.7260 | kinesin | **Flagellum-puncta** | 5 | 1.32 |
| Tb927.9.9690 | Hypothetical | **Flagellum, Tip-enriched** | TrypTag<sup>1</sup> | 1.22 |
| Tb927.7.3090 | Galactose Oxidase-central domain --containing | **Not Flagellar** | TrypTag<sup>1</sup> | 1.17 |
| Tb927.5.2410 | kinesin | **Flagellum, Tip-enriched** | TrypTag<sup>1+6</sup> | 1.16 |

\*Localization defined as: Tip = the prominent or sole location is flagellum tip. Flagellum, Tip-enriched = in the flagellum and enriched at tip. Tip-enriched = enriched at the flagellum tip, but also present outside the flagellum.
<sup>1</sup>Dean, S., J.D. Sunter, and R.J. Wheeler, TrypTag.org: A Trypanosome Genome-wide Protein Localisation Resource. Trends Parasitol, 2017. 33(2): p. 80-82.
<sup>2</sup>Subota, I., *et al.*, Proteomic analysis of intact flagella of procyclic *Trypanosoma brucei* cells identifies novel flagellar proteins with unique sub-localization and dynamics. Mol Cell Proteomics, 2014. 13(7): p. 1769-86.
<sup>3</sup>Liu, W., *et al.*, Expression and cellular localisation of calpain-like proteins in *Trypanosoma brucei*. Mol Biochem Parasitol, 2010. 169(1): p. 20-6.
<sup>4</sup>Saada, E.A., *et al.*, Insect stage-specific receptor adenylate cyclases are localized to distinct subdomains of the *Trypanosoma brucei* Flagellar membrane. Eukaryot Cell, 2014. 13(8): p. 1064-76.
<sup>5</sup>Demonchy, R., *et al.*, Kinesin 9 family members perform separate functions in the trypanosome flagellum. J Cell Biol, 2009. 187(5): p. 615-22.
<sup>6</sup>An, T. and Z. Li, An orphan kinesin controls trypanosome morphology transitions by targeting FLAM3 to the flagellum. PLoS Pathog, 2018. 14(5): p. e1007101.

Prevalence of adenylate cyclases within the AC1s proximity proteome supports the idea that the dataset is enriched for tip proteins, because all *T. brucei* adenylate cyclases studied to date are flagellar and, in procyclics, many are enriched at the flagellum tip [29]. AC2 is localized all along the flagellum [29], yet it is found in the AC1s proximity proteome, perhaps due to the fact that AC2 and AC1 dimerize and share ∼90% amino acid sequence identity [29].

Among the 21 non-AC proteins in the AC1s proximity proteome, 20 have independent data on localization [43]. Four of these have previously been published as being flagellum tip-specific (FLAM8 and CALP1.3), flagellum-specific and tip-enriched (KIN-E), or located throughout the cell but also found in the flagellum tip (CALP7.2) [34, 53, 54]. For the remaining 16 proteins we assessed localization by referencing the TrypTag database [43] and/or epitope tagging directly. We find that half of these 16 proteins are either tip-specific or enriched at the flagellum tip while also being located elsewhere in the cell (Fig. 7 and Table 2). Notably, most of the proteins that did not exhibit tip localization were among the least enriched in the AC1s versus FS179s samples (Table 2). Among 11 proteins enriched >2-fold in AC1s versus FS179s and having localization data, 9 are enriched in the flagellum tip. Therefore, the AC1s proximity proteome is enriched for flagellum tip proteins.

**Figure 7.**
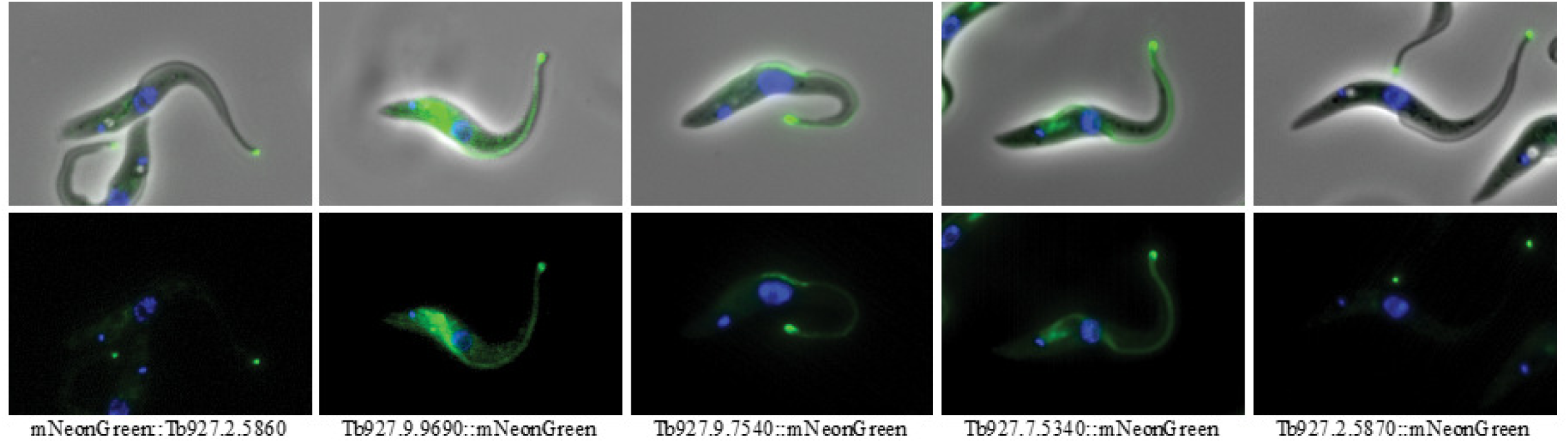
AC1s proximity proteome identifies tip proteins. Fluorescence microscopy of trypanosomes expressing the indicated protein tagged with neon green (Green). Samples are stained with DAPI (Blue). Top panel shows fluorescence plus phase contrast merged images. Bottom panel shows fluorescence image.

## DISCUSSION

The protein composition of flagellum subdomains in *T. brucei* is a knowledge gap in understanding the biology of these pathogens. To overcome this, we implemented APEX2 proximity proteomics. Our results demonstrate that APEX-based proximity labeling is effective in *T. brucei* and is capable of resolving flagellum subdomains even if they are not separated by physical barriers. Use of the APEX system has allowed us to define a soluble flagellum tip proteome that indicates the tip is enriched for signaling proteins. While this tip proteome is likely incomplete, our work represents an important step in defining protein composition of flagellum subdomains and other trypanosome cellular compartments that cannot themselves be purified.

One major advantage of proximity labeling-based proteomics versus other proteomic approaches to define organelle protein composition is that it allows for isolation of proteins of interest from crude cell lysates in a simple, one-step purification, without the need to purify the organelle. Prior proteomic analyses of the *T. brucei* flagellum have required purification of the flagellum away from the cell body [34]. This is problematic, because the flagellum in *T. brucei* is laterally connected to the cell body along most of its length. Therefore, purification approaches have employed genetic manipulation to remove lateral connections, followed by sonication or shearing to detach the flagellum at its base [32, 34] or have employed detergent extraction followed by selective depolymerization of subpellicular microtubules while leaving axoneme microtubules intact [30]. The latter approach does not allow for identification of flagellum matrix or membrane proteins that are detergent-soluble. In both cases, flagellum detachment is followed by centrifugation to separate flagellum fractions from cell bodies and solubilized material, then electron microscopy to evaluate sample quality. The APEX approach eliminates the need for subcellular fractionation and evaluation of the purified flagellum sample, although it can be coupled with fractionation to distinguish detergent-soluble from insoluble proteins, as done in our case. APEX labeling also enables easy detection of detergent-soluble and insoluble proteins from the same cells. A disadvantage of proximity labeling-based proteomics is that owing to the requirement for proteins to be close to the APEX-tagged bait, one might miss proteins that are part of the organelle in question, but distant from the bait protein. For this reason, efforts to define a comprehensive, whole-organelle proteome should employ multiple independent approaches.

Our ability to distinguish protein profiles of AC1s and FS179s samples (Fig 6A) illustrates the capacity of APEX to distinguish between subcellular regions that are not separated by a physical barrier. Although this capacity was shown previously in mammalian cells for the cilium compartment versus the cytoplasm [44] and for distinct locations in the cytoplasm [55], our studies now demonstrate this is also possible within the spatially-restricted volume of the flagellum. The eukaryotic flagellum is comprised of specific subdomains, each with specialized functions and protein composition [29]. The distal tip of the flagellum in many organisms, for example, is important for transduction of extracellular signals, flagellum length regulation and cell-cell adhesion [10, 11, 45–51]. The transition zone at the flagellum’s proximal end has specialized functions controlling access into and out of the flagellar compartment [16, 56]. Even dyneins are distributed differentially from proximal to distal ends of the axoneme [57]. APEX labeling has previously been used to identify proteins throughout the cilium in mammalian cells [44]. To our knowledge however, our studies are the first to extend this system to distinguishing protein composition of specific flagellum subdomains, thus providing a powerful addition to tools available for dissecting flagellum function and mechanisms of ciliary compartmentation.

Prior work has provided important advances by determining protein composition of two specific subdomains within the *T. brucei* flagellum, the transition zone [16] and tip [58] of the detergent-insoluble axoneme, the latter including an “axonemal capping structure” (ACS) and the *T. brucei*-specific flagellar connector (FC). Those studies developed a novel method termed “structure immunoprecipitation” (SIP), in which detergent-insoluble axonemes are prepared, then fragmented into small pieces and immunoprecipitated using antibody to a marker protein localized to the domain of interest. Independent localization, biochemical and functional analysis were then used to define a corresponding transition zone, ACS and FC proteomes. Given that the FC and ACS are at the tip of the flagellum, we compared their proteomes to the AC1s proximity proteome. We did not find any overlap, as might be expected because the FC and ACS analyses were restricted to detergent-insoluble components, while the AC1s proximity proteome uniquely identifies detergent-soluble proteins. Our work therefore complements and extends earlier proteome analyses of *T. brucei* flagellum sub-domains and expands our knowledge of proteins that mediate flagellum tip-specific functions.

The AC1s proximity proteome is enriched for known and previously unknown flagellum tip proteins (Fig. 7 and Table 2). Function for many of these proteins remains to be determined but features and properties in some cases indicate a role in signal transduction. One, previously identified as “cAMP response protein 3” (CARP3), is a candidate cAMP effector [59] and over half of the proteins are adenylate cyclases. A role for the flagellum tip in cAMP signaling has been previously demonstrated [10, 11] and flagellar cAMP signaling is required for the *T. brucei* transmission cycle in the tsetse fly [12]. Another protein group well-represented in the AC1s proximity proteome is calpain-like proteins. Calpains function in Ca^++^ signal transduction, although the calpain-like proteins in the AC1s proximity proteome possess only a subset of the domains typically seen in more classical calpains. Interestingly, the flagellar tip kinesin KIN-E [53] contains a domain III-like domain typically found in calpains and thought to function in lipid binding [60]. The AC1s proximity proteome includes four proteins annotated as having homology to a serine-rich adhesin protein from bacteria [61]. Whether these function as adhesins in the flagellum is unknown, but adhesion functions could contribute to attachment of the axoneme to the flagellar membrane or in attachment functions of the FC [58].

The DRC1p proximity proteome showed substantial overlap with earlier proteome analyses of purified flagellum skeletons [16, 30, 31, 34], while also including 350 proteins not found in those prior studies. Independent data on 241 of these 350 proteins support flagellum association for 71%, indicating protein composition of the *T. brucei* flagellum is more complex than suggested from earlier studies. Many proteins uncovered in prior flagellum proteome analyses were not found in the DRC1p proximity proteome. There are several potential explanations for this. First, some earlier studies include detergent-soluble matrix and membrane proteins [32, 34], which are not expected in the DRC1p proximity proteome. Second, the threshold for inclusion in the DRC1p dataset is high, as proteins must be present in all four DRC1p replicates and enriched over negative controls, while some of the earlier studies examined a limited number of replicates [30], or lacked extensive negative controls [32]. Third, each approach will contain false positives and false negatives. Lastly, proteins that are far away from DRC1 might not be biotinylated in our analysis. We recognize that several bona fide axonemal proteins were identified in WT, negative control samples processed in parallel to DRC1p samples. We do not presently know the reason for this, but the problem was less evident in the analysis of detergent-soluble AC1s samples, so may relate to residual insolubility of the axoneme, or high abundance of axoneme proteins, which may be present at thousands of copies per axoneme.

To our knowledge, this is the first work to demonstrate APEX2 proximity proteomics in a pathogenic eukaryote. This is an important point, because presence of systems for removing reactive oxygen species in any given organism may limit suitability of APEX-based labeling [36]. For example, *Plasmodium spp.* rely on a series of redundant glutathione- and thioredoxin-dependent reactions to remove H_2_O_2_ and maintain redox equilibrium [62]. BioID [63] is an alternative proximity-labeling method that has been used widely, including several applications in *T. brucei* [64] and other parasites [65]. BioID and APEX methods are complementary, with APEX offering the added advantage of capacity for doing time course analyses. Another advantage is that APEX catalyzes oxidation of 3,3’-diaminobenzidine (DAB), which polymerizes and can be visualized by electron microscopy to determine precise subcellular location of the APEX-tagged protein [66]. Therefore, our studies expand the capacity in *T. brucei* for proximity-labeling, which is an increasingly important tool in post-genomic efforts to define protein location and function in pathogenic organisms [67].

## MATERIALS AND METHODS

### *Trypanosoma brucei* culture

Procyclic *Trypanosoma brucei brucei* (strain 29-13) [68] was used as control and to generate all the APEX2-tagged cell lines. Cells were cultivated in SM media supplemented with 10% heat inactivated fetal bovine serum and incubated at 28°C with 5% CO2. Selection for transformants was done using 1 μg/mL of puromycin.

### In situ Tagging

APEX2-tagged cell lines were generated by *in situ* tagging [69]. In each case, cells were transfected with a cassette containing the APEX2-NES tag [35] followed by a puromycin resistance marker and flanked on the 5’ end by the 3’ end of the target gene ORF and on the 3’ end by the target gene 3’ UTR. For AC1 (Tb927.11.17040) and FS179 (Tb927.10.2880), the AC1-HA [32] and FS179-HA [29] *in situ* tagging vectors were modified by inserting the APEX2-NES tag in-frame between the target gene ORF and HAx3 tag to generate the final APEX2 tagging cassettes. For DRC1 (Tb927.10.7880), the last 592 bp of the ORF were PCR-amplified from genomic DNA and cloned upstream of the HAx3 tag in the pMOTag2H [69] vector backbone. Similarly, the first 404 bp of the DRC1 3’-UTR were PCR-amplified from genomic DNA and cloned downstream of the puromycin resistance marker. The APEX2-NES tag was then inserted between the target gene ORF and HAx3 tag as described above. All sequences were verified by DNA sequencing. Tagging cassettes were excised from the pMOTag backbone and gel purified. Trypanosome cells were transfected by electroporation and selected with puromycin as described [39].

### Biotinylation

Biotinylation was done using a modified version of [36]. Briefly, cells were harvested by centrifugation for 10 min at 1,200 x g, then resuspended at 2 × 10^7^ cells/mL in growth medium supplemented with 5 mM biotin-phenol (biotin tyramide, Acros Organics). After a one-hour incubation, cells were treated with 1 mM H_2_O_2_ for one minute. To quench unreacted hydrogen peroxide, an equal volume of 2x quenching buffer (10 mM Trolox and 20 mM L-Ascorbic Acid Sodium Salt in PBS, pH 7.2) was added, cells were harvested by centrifugation and two additional washes were made with 1x quenching buffer.

### Immunofluorescence

Cells were washed once in PBS and fixed by addition of paraformaldehyde to 0.1% for 5 min on ice. Fixed cells were washed once in PBS and air-dried onto coverslips. The coverslips were incubated for 10 min in −20°C methanol followed by 10 min in −20°C acetone, and then air-dried. Cells were re-hydrated for 15 min in PBS and blocked overnight in blocking solution (PBS + 5% bovine serum albumin [BSA] + 5% normal donkey serum). Coverslips were incubated with Streptavidin coupled to Alexa 594 (Life Technologies) and anti-PFR [29] diluted in blocking solution for 1.5 h. After three washes in PBS + 0.05% Tween-20 for 10 min each, coverslips were incubated with donkey anti-rabbit IgG coupled to Alexa 488 (Invitrogen) in blocking solution for 1.5 h. After three washes in PBS + 0.05% Tween-20 for 10 min each, cells were fixed with 4% paraformaldehyde for 5 min. Coverslips were washed three times in PBS + 0.05% Tween-20 and one time in PBS for 10 min each. Cells were mounted with Vectashield containing DAPI (Vector). Images were acquired using a Zeiss Axioskop II compound microscope and processed using Axiovision (Zeiss, Inc., Jena, Germany) and Adobe Photoshop (Adobe Systems, Inc., Mountain View, CA).

### Fractionation of Whole Cells and Purification of Biotinylated Proteins

For purification of biotinylated proteins, 6 × 10^8^ cells were washed once in PBS and lysed in lysis buffer: PEME buffer (100 mM PIPES • 1.5 Na, 2 mM EGTA, 1 mM MgSO_4_ • 7 H_2_O, and 0.1 mM EDTA-Na_2_ • 2 H_2_O) + 0.5% Nonidet P-40 (NP40) + EDTA free protease inhibitors (Sigma), for 10 min on ice. Lysates were centrifuged for 8 min at 2,500 x g at room temperature (RT) to separate the NP40-soluble supernatant and the insoluble pellet. The pellet was boiled for 5 min in lysis buffer + 1% SDS, centrifuged for 3 min at 21,000 x g at RT to remove insoluble debris and SDS supernatant collected.

Capture of biotinylated proteins from the NP40-soluble and SDS supernatants on streptavidin beads was done essentially as described [32]. Cell fractions were incubated with 120 µL streptavidin beads (GE Healthcare) overnight at 4°C with gentle agitation. Biotinylated proteins bound to Streptavidin beads were separated from the unbound molecules by centrifugation. Beads were washed once with lysis buffer at 4°C, whereas the rest of the washes were made at RT as follows: once in buffer A (8 M urea, 200 mM NaCl, 2% SDS, and 100 mM Tris), once in buffer B (8 M urea, 1.2 M NaCl, 0.2% SDS, 100 mM Tris, 10% ethanol, and 10% isopropanol), once in buffer C (8 M urea, 200 mM NaCl, 0.2% SDS, 100 mM Tris, 10% ethanol, and 10% isopropanol), and 5 times in buffer D (8 M urea, and 100 mM Tris); pH of all wash buffers was 8.

### Shotgun Proteomics

Shotgun proteomics was done based on [32]. Streptavidin-bound proteins were digested on beads by the sequential addition of lys-C and trypsin protease. Peptide samples were fractionated online using reversed phase chromatography followed by tandem mass spectrometry analysis on a Thermofisher Q-Exactive™ mass spectrometer (ThermoFisher). Data analysis was performed using the IP2 suite of algorithms (Integrated Proteomics Applications). Briefly, RawXtract (version 1.8) was used to extract peaklist information from Xcalibur-generated RAW files. Database searching of the MS/MS spectra was performed using the ProLuCID algorithm (version 1.0) and a user assembled database consisting of all protein entries from the TriTrypDB for *T. brucei* strain 927 (version 7.0). Other database search parameters included: 1) precursor ion mass tolerance of 10 ppm, 2) fragment ion mass tolerance of 10ppm, 3) only peptides with fully tryptic ends were considered candidate peptides in the search with no consideration of missed cleavages, and 4) static modification of 57.02146 on cysteine residues.

Peptide identifications were organized and filtered using the DTASelect algorithm which uses a linear discriminate analysis to identify peptide scoring thresholds that yield a peptide-level false discovery rate of less than <1.8% as estimated using a decoy database approach. Proteins were considered present in the analysis if they were identified by two or more peptides using the <1.8% peptide-level false discovery rate.

For principal component analysis (PCA) and comparison of proteins identified in each sample, proteomics data were parsed using the IP2 Integrated Proteomics Pipeline Ver. 5.1.2. The output from the Protein Identification STAT Compare tool (IDSTAT_COMPARE) in IP2 was processed using a custom R-based web app (https://uclaproteomics.shinyapps.io/iscviewer/) to generate the PCA graphs. For proteins identified in different samples, the ID_COMPARE output was exported to Excel to generate mass spectrometry data analysis tables (Supplemental Tables S4 and S5).

For the DRC1p proximity proteome, data were from three independent experiments each for DRC1-APEX2 and 29-13 (WT) pellet samples. In one case each, the sample was split into two aliquots and shotgun proteomics was done on both in parallel, giving a total of four replicates each for DRC1-APEX2 and WT samples. Ratios of average normalized spectra were used to determine inclusion in the DRC1p proximity proteome as described in the text. Of 739 proteins identified, 42 redundant GeneIDs were removed for a total proximity proteome of 697 (Table S1 and S4). For the AC1s proximity proteome, data were from four independent experiments for 29-13 samples (0313, 0410, 0629-A and 0629-B) and two independent experiments each for AC1 (0313, 0410) and FS179 (0629-A and O629-B). For the 0313 and 0410 experiments, each sample was split into two aliquots and shotgun proteomics done in parallel on each, for a total of six replicates for WT and four replicates for AC1. Ratios of average normalized spectra were used to determine inclusion in the AC1s proximity proteome as described in the text.

### Bioinformatics

To identify human homologues, we developed an algorithm to automatically return reciprocal best blast hits from the large lists of proteins produced by shotgun proteomics. The algorithm was implemented in python using the biopython library (https://academic.oup.com/bioinformatics/article/25/11/1422/330687) and works as follows. Individual sequences were parsed sequentially and used as a query for BLASTp to find a list of similar sequences with an e-value threshold of 0.1 from a database of human protein sequences retrieved from NCBI. The top three most similar sequences were then used as query sequences for a subsequent BLASTp call against a database of *T. brucei* protein sequences. If the original sequence was found in the top three from this call then the query was then returned as a homologue. The python code for performing this can be found at the following URL: https://github.com/marcusgj13/Reciprocal-BB. Comparison to published *T. brucei* flagellar proteomes [30, 32, 34] was done using the search tools in the TriTryp Genome Database (https://tritrypdb.org/tritrypdb/) [61]. Word clouds (Figures 3, 6 and S2) were obtained using the GO Enrichment tool of TriTrypDB, using *T. brucei brucei* TREU927 and a 0.05 P-Value cutoff.

To assess protein localization as determined by TrypTag [43], proteins in the DRC1p proximity proteome dataset were cross-referenced with TrypTag to look at matches vs mis-matches. The GO terms searched for were: flagellum, transition zone, flagellar cytoplasm, pro-basal body, basal body, flagellar membrane, flagella connector, paraflagellar rod, flagellum attachment zone, intraflagellar transport particle, hook complex, flagellar tip, and axoneme. For the full DRC1p proximity proteome, the number of genes that were searched for was 739. The number of genes for which there is TrypTag localization data for at least one terminus was 677. The number of TrypTag-tagged genes that matched a flagellum GO term were 509. The number of TrypTag-tagged genes that DID NOT match a flagellum GO term was 168. Therefore of 677 proteins with TrypTag localization data, 75% have a TrypTag localization that matches the query GO terms. For DRC1p proximity proteome proteins not found in earlier proteome analyses, the number of query genes searched for was 350. The number of query genes for which there is TrypTag localization data for at least one terminus was 337. The number of query genes where TrypTag localization MATCHES the query GO term was 241. The number of query genes where TrypTag localization MIS-MATCHES the query GO term was 96. Therefore, of 337 proteins with TrypTag localization data, 71% have a TrypTag localization that MATCHES the query GO terms.

### Data availability

The python code used to identify human homologues was deposited in GitHub, and it can be found at the following URL: https://github.com/marcusgj13/Reciprocal-BB. The code, explanation of it, as well as instructions to install and run the program can be found there.

## ACKNOWLEDGMENTS

We thank Vincent Tran for help with molecular cloning. Funding was provided by National Institutes of Health grants AI052348 and AI142544 to K.H., and GM089778 to J.A.W. The authors would like to thank the TrypTag consortium, which is supported by the Wellcome Trust [108445/Z/15/Z], for providing data which enabled part of this work. D.E.V.R. was recipient of a Fulbright scholarship trough the United States-Mexico Commission for Educational and Cultural Exchange (COMEXUS), a postdoctoral fellowship of the Mexican National Council of Science and Technology (CONACyT) (CVU #325790), and a postdoctoral fellowship of the University of California Institute for Mexico and the United States (UC MEXUS), in conjunction with CONACyT. The funders did not have any role on the study design, data collection and interpretation, nor the decision to submit the work for publication.

## Legends

**Supplemental Figure S1. Growth curves of APEX2-tagged cell lines.**

Comparison of doubling times of 29-13 (parental cell line), AC1-APEX2, DRC1-APEX2, and FS179-APEX2 cell lines in suspension culture over the course of 8 days. Cell densities were adjusted daily to 1e6 cells/mL in order to insure logarithmic growth.

**Supplemental Figure S2. Word cloud representing GO analysis for Cellular Component (A) and Biological Process (B) of the proteins identified in the DRC1s proximity proteome.**

**Supplemental Figure S3. FAZ proteins identified on the DRC1p proximity proteome.**

Panel A is a schematic of the FAZ. Panel B indicates the FAZ zone to which each protein has been localized, as well as the ratio of spectra identified in the DRC1p versus 2913p sample. FAZ schematic and zone locations are based on Sunter, J.D. and K. Gull, The Flagellum Attachment Zone: ‘The Cellular Ruler’ of Trypanosome Morphology. <u>Trends Parasitol</u>, 2016. 32(4): p. 309-324.

